# Label-free imaging to track reprogramming of human somatic cells

**DOI:** 10.1101/2021.12.08.471827

**Authors:** Kaivalya Molugu, Giovanni A. Battistini, Tiffany M. Heaster, Jacob Rouw, Emmanuel C. Guzman, Melissa C. Skala, Krishanu Saha

## Abstract

The process of reprogramming patient samples to human induced pluripotent stem cells (iPSCs) is stochastic, asynchronous, and inefficient leading to a heterogeneous population of cells. Here, we track the reprogramming status of single patient-derived cells during reprogramming with label-free live-cell imaging of cellular metabolism and nuclear morphometry to identify high-quality iPSCs. Erythroid progenitor cells (EPCs) isolated from human peripheral blood showed distinct patterns of autofluorescence lifetime for the reduced form of nicotinamide adenine dinucleotide (phosphate) [NAD(P)H] and flavin adenine dinucleotide (FAD) during reprogramming. Random forest models classified starting EPCs, partially-reprogrammed intermediate cells, and iPSCs with ∼95% accuracy. Reprogramming trajectories resolved at the single cell level indicated significant reprogramming heterogeneity along different branches of cell state. This combination of micropatterning, autofluorescence imaging, and machine learning provides a unique non-destructive method to assess the quality of iPSCs in real-time for various applications in regenerative medicine, cell therapy biomanufacturing, and disease modeling.

## Introduction

The derivation of patient-specific induced pluripotent stem cells (iPSCs) from their somatic cells via reprogramming generates a unique self-renewing cell source for disease modeling, drug discovery, toxicology, and personalized cell therapies^1–3^. These cells carry the genome of the patient, facilitating elucidation of the genetic causes of disease, and are immunologically matched to the patient, facilitating the engraftment of any cell therapies developed from these cells^4–6^.

With several clinical trials planned and underway^7^, there has been significant progress in developing iPSC-based cell therapies in recent years. However, several challenges remain in the field^8^. First, the derivation of high-quality iPSCs must be efficient, rapid, and cost-effective to ensure that patients receive their treatments in a timely fashion for autologous iPSC-derived products. Second, iPSC-derived cell therapies for both allogeneic and autologous strategies require scalable and standardized manufacturing processes to overcome the inconsistencies arising from variability in human material sources, reagents, delivery of the reprogramming factors, microenvironmental fluctuations, or inherent stochasticity in epigenetic processes underpinning reprogramming^9^. Typical assays currently used for quality control of GMP (good manufacturing practices)-grade iPSCs include testing for cell line identity (STR analysis, SNP analysis, genomic sequencing), genomic instability (G-banding, chromosomal microarray, Nanostring technology), pluripotency (marker expression analysis via flow cytometry or immunochemistry, embryoid body analysis, teratoma assays, Pluritest™, TaqMan Scorecard™ Assay) and residual expression of reprogramming factors (PCR or immunochemistry)^10–13^. Each of these methods can be low-throughput, labor-intensive, time-consuming, and require destructive processing.

Using non-destructive strategies, recent studies have indicated that automated machine learning can be used to identify cell structures from label-free brightfield images that cannot be manually identified^14–18^. However, such automated methods to identify iPSCs based on cellular morphology have had limited success in the field^19–21^. Deep learning has recently been developed to analyze monoclonal cell cultures of iPSC^22^, however reprogramming cultures involve a higher number of cell fate transitions that have yet to be analyzed through deep learning pipelines. Hence, in complex cultures, like those in reprogramming, new standardized platforms with robust analytical methods for identifying high-quality iPSCs are still needed.

Our strategy to identify iPSCs and other cells during reprogramming exploits metabolic and nuclear changes during reprogramming. Somatic cells primarily utilize mitochondrial oxidative phosphorylation (OXPHOS) to support cell proliferation^23^. However, pluripotent stem cells favor glycolysis, in a manner reminiscent of the Warburg effect in cancer cells^23,24^. During reprogramming, somatic cells thus undergo a metabolic shift from OXPHOS to glycolysis^25,26^, triggered by a transient OXPHOS burst, resulting in initiation and progression of reprogramming to iPSCs^27–29^. Recent evidence also indicates that this metabolic shift occurs prior to changes in gene expression and that the modulation of glycolytic metabolism or OXPHOS alters reprogramming efficiency^24,30,31^. High-resolution imaging of reprogramming cells has also identified that nuclear geometry is dramatically altered during reprogramming^32–34^. Therefore, simultaneous monitoring metabolic and nuclear changes during reprogramming could reveal insights into reprogramming and subsequent identification of iPSCs.

Optical Metabolic Imaging (OMI), a non-invasive and label-free two-photon microscopy technique that provides dynamic measurements of cellular metabolism at a single-cell level. OMI is based on the endogenous fluorescence of metabolic coenzymes, NADH, and FAD^35^, which are both used across several cellular metabolic processes. NADH and NADPH have overlapping fluorescence properties and are collectively referred to as NAD(P)H^36^. The optical redox ratio, defined as the ratio of NAD(P)H intensity to total NAD(P)H and FAD intensity, provides a measure of the relative oxidation-reduction state of the cell [Inad(p)h/(Inad(p)h+Ifad)]^37,38^. Fluorescence lifetime imaging microscopy (FLIM) of NAD(P)H and FAD provides additional information specific to protein binding activity. The two-component decays of NAD(P)H and FAD measure the short (τ_1_) and long (τ_2_) fluorescence lifetimes that correspond to the free or bound states of these coenzymes^39–41^, along with fractional contributions of short (α_1_) and long (α_2_) lifetimes. Since NAD(P)H and FAD are found predominantly in the cytoplasm, the lack of fluorescence signal in images can also be used to identify cell borders and nuclei^42^. Thus, OMI provides multiple readouts for cell metabolism and nuclear morphometry to track metabolic and nuclear changes of cells undergoing reprogramming.

Here, we address some of the challenges associated with the biomanufacturing of iPSCs by developing a microcontact printed (μCP) platform^34,43,44^ to non-invasively monitor metabolic and nuclear changes over 22 days of reprogramming of human EPCs to iPSCs. With this study, we demonstrated that OMI is sensitive to the metabolic and nuclear differences during reprogramming, performed accurate classification of reprogramming status of cells using machine learning models, and subsequently built single-cell reprogramming trajectories^45^. Our label-free, non-destructive, rapid, scalable method to track reprogramming provides novel insights at the single cell level into human cell reprogramming and could enable the development of new technologies for biomanufacturing high-quality iPSCs.

## Results

### Metabolic imaging during reprogramming on patterned substrates

We first designed a microcontact printed (μCP) substrate to spatially control the adhesion of EPCs undergoing reprogramming^34,43,46^. The μCP substrate is formed by coating 300μm radius circular regions, referred to as μFeatures, with Matrigel on a 35mm ibiTreat dish that allows for cell adhesion. The remaining regions of the dish are then backfilled with polycationic graft copolymer, PLL-g-PEG, that resists protein adsorption and prevents cell adhesion in these regions^47,48^ (**Fig. S1a**). The ibiTreat dishes are made of gas-permeable material, enabling maintenance of carbon dioxide or oxygen exchange during cell culture and have high optical quality. These properties make the dishes suitable for two-photon microscopy during reprogramming. To verify proper coating of the circular μFeature regions, we immunostained for laminin, a major component of Matrigel^49^. Fluorescence imaging showed laminin consistently within the circular μFeatures indicating uniform patterning of Matrigel (**Fig. 1a**). We next assessed the ability of the μCP substrates to enable cell attachment by seeding two different cell types i.e., human dermal fibroblasts (HDFs) and H9 human embryonic stem cells (H9 ESCs). We observed that both HDFs and ESCs remained viable, attached, and confined to the circular μFeatures indicating that the μCP substrates enable spatial control of cell adhesion (**Fig. S1b**).

**Fig. 1.**
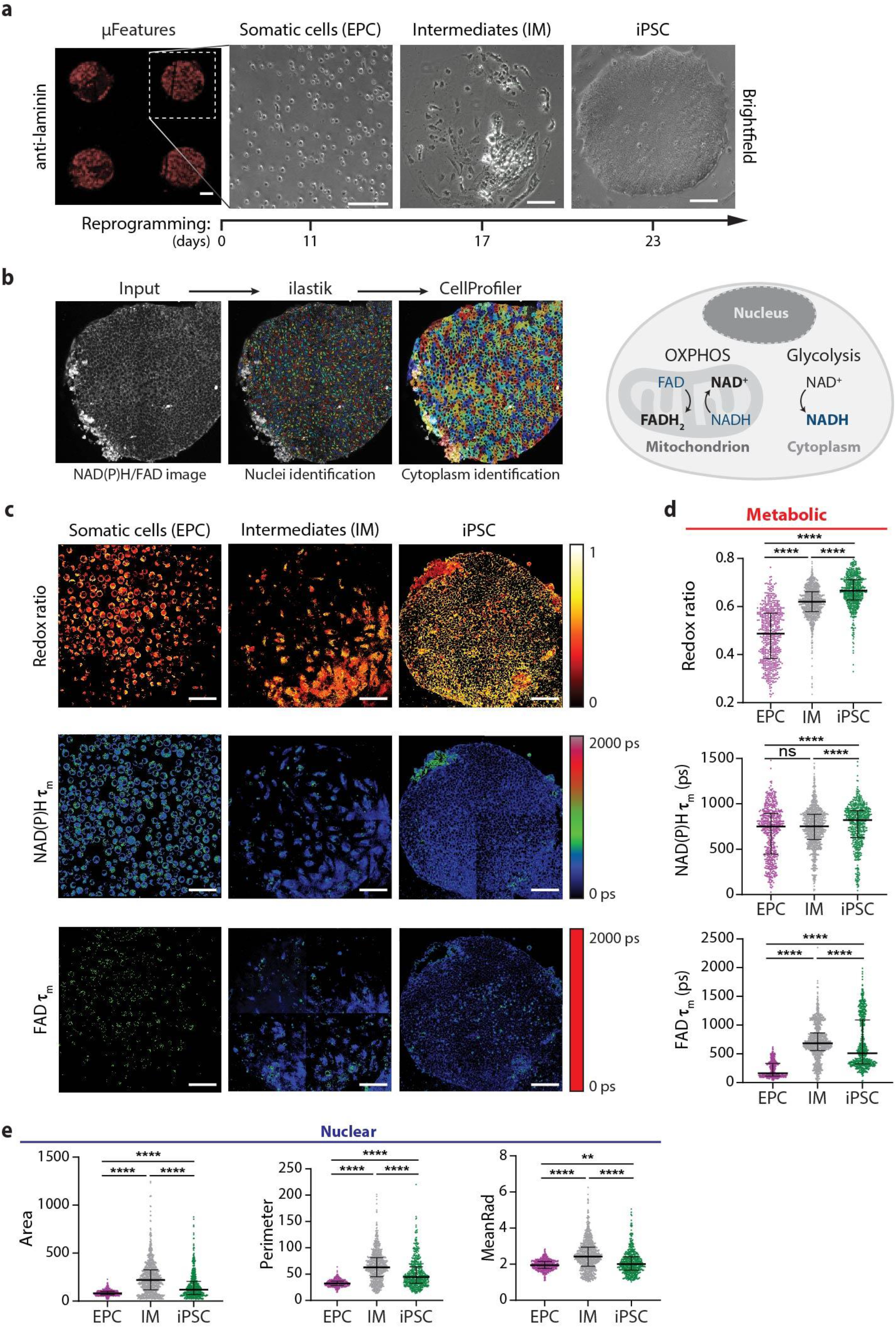
NAD(P)H and FAD autofluorescence imaging reveal metabolic differences during reprogramming. **a) Left:** Matrigel-coated μFeatures on the ibiTreat dish visualized with an anti-laminin antibody (red) show good fidelity in the transfer from the Matrigel-coated PDMS mold. Scale bar, 100 μm. **Right:** Representative images of the progression of erythroid progenitor cells (EPCs) on a single circular μFeature (300 μm radius) through a reprogramming time course. **b) Left:** Image analysis pipeline to identify metabolic and nuclear parameters using ilastik and CellProfiler software. **Right:** Schematic representation of cell metabolism with NADH and FAD highlighted as the fluorescent molecules in the diagram, and molecules in bold indicate the net direction of the reaction. **c)** Representative optical redox ratio, NAD(P)H τ_m_ and FAD τ_m_ images (3 μFeatures selected from 36 μFeatures acquired from 3 different donors) for EPC, IM, and iPSC. Color bars are indicated on the right. Scale bar, 100 μm. **d)** Single-cell quantitative analysis of metabolic parameters: optical redox ratio, NAD(P)H τ_m_, FAD τ_m_, FAD α_1_; and nuclear parameters: area, perimeter, mean radius (*n* = 561, 990, 586 for EPC, IM, and iPSC respectively). Data are presented as median with interquartile range for each cell type. Statistical significance was determined by one-way analysis of variance (ANOVA) using the Kruskal-Wallis test for multiple comparisons; ns = p ≥0.05, * for p <0.05, ** for p <0.01, *** for p <0.001, **** for p <0.0001.

Next, we isolated peripheral blood mononuclear cells (PBMCs) from peripheral blood of healthy human donors and further enriched them for EPCs prior to the delivery of reprogramming factors. We examined the enrichment of EPCs by flow cytometry with erythroid cell surface marker CD71^50^. Flow cytometry confirmed the presence of enriched EPCs with flow cytometry showing that >98% of the cells expressed CD71 on day 10 of PBMC culture (**Fig. S1c**).

To initiate reprogramming, we electroporated the EPCs with four episomal reprogramming plasmids^51,52^ — encoding Oct4, shRNA knockdown of p53, Sox2, Klf4, L-Myc, Lin28, and miR302-367 cluster — and seeded them onto μCP substrates. We assessed the ability of the μCP substrates to sustain long-term reprogramming studies by performing high-content imaging to track individual μFeatures (>30 μFeatures per 35mm dish) longitudinally at multiple timepoints over the ∼3-week of reprogramming. Day 22 was picked as the reprogramming endpoint because there were several iPSC colonies at this timepoint without significant outgrowth within a μFeature. While the starting EPCs are non-adherent, cells in the middle of reprogramming, termed intermediate cells (IMs), and endpoint iPSCs adhere to the circular μFeatures within the μCP substrates (**Fig. 1a**) indicating that μCP substrates can support the full reprogramming of EPCs. Overall, the μCP platform provides unique spatial control over reprogramming cells and enables high-content quantitative imaging of reprogramming.

### OMI reveals distinct metabolic changes during reprogramming

Metabolic state plays an important role in regulating reprogramming and pluripotency of iPSCs^53–57^ and can be non-invasively monitored via OMI. NAD(P)H is an electron donor and FAD is an electron acceptor. Both are present in all cells as coenzymes and provide energy for metabolic reactions. For example, glycolysis in the cytoplasm generates NADH and pyruvate, while OXPHOS consumes NADH and produces FAD (**Fig. 1a**). Autofluorescence imaging of NAD(P)H and FAD is thus dynamically responsive to the oxidation-reduction state of a cell and is influenced by many reactions^35,58^.

We tracked the autofluorescence dynamics of NAD(P)H and FAD by performing OMI on μCP substrates at different time points during EPC reprogramming. In these images, the nucleus remains dark as NAD(P)H is primarily located in cytosol and mitochondria, and FAD is primarily located in mitochondria. The NAD(P)H images were used as inputs for ilastik software^59^ to identify the nuclei. The identified nuclei were then used as an input for high-content CellProfiler software^60^ pipeline to segment the cytoplasm, and measure various metabolic and nuclear parameters (**Fig. 1b**). Altogether, 11 metabolic parameters (NAD(P)H intensity, Inad(p)h; NAD(P)H α_1_; NAD(P)H τ_1_; NAD(P)H τ_2_; NAD(P)H mean lifetime, τ_m_ = α_1_τ_1_ + α_2_τ_2_, FAD intensity, Ifad; FAD α_1_; FAD τ_1_; FAD τ_2_; FAD τ_m_; optical redox ratio, Inad(p)h / [Inad(p)h + Ifad]), and 8 nuclear parameters^34^ (area; perimeter; mean radius, MeanRad; nuclear shape index, NSI; solidity; extent; number of neighbors, #Neigh; distance to closest neighbor, 1stNeigh) were measured by the analysis pipeline (**Fig. S1d**). By fixing the cultures at these time points, we verified the cell type by immunofluorescence labeling: EPCs (CD71^+^, Nanog^-^), IMs (CD71^-^, Nanog^-^), and iPSCs (CD71^-^, Nanog^+^). NAD(P)H and FAD autofluorescence imaging revealed metabolic differences between starting EPCs, intermediates (IM), and iPSCs (**Fig. S2, S3**).

We observed a significant increase in the optical redox ratio (iPSC>IM>EPC) during reprogramming (**Fig. 1c**), indicating that individual erythroid progenitor cells are more oxidized than individual IMs and iPSCs (**Fig. 1d**). Additionally, we noted that patterned IMs and iPSCs have significantly higher optical redox ratios as compared to their non-patterned counterparts (**Fig. S2k**). This observation is consistent with previous studies which show that mechanical cues can regulate their relative use of glycolysis^61–64^.

Next, we observed that NAD(P)H and FAD lifetime components undergo biphasic changes during the progress of reprogramming. For both phases, FAD lifetime components undergo a more significant change relative to the NAD(P)H components (**Fig. 1d, Fig. S2a-j**). On average, the fraction of protein-bound FAD (FAD α_1_) first decreases from its levels in EPCs to those in IMs and then increases during the IM to iPSC level, which could be reflective of the OXPHOS burst^27– 29^ (**Fig. S2h**). FAD τ_m_ (**Fig. 1d**) is inversely related to FAD α_1_ and therefore undergoes a biphasic change that is opposite to that of FAD α_1_. Similar biphasic changes occur in nuclear parameters during reprogramming, which is consistent with our previous study^34^ (**Fig. 1e, Fig. S3**).

We compared these measurements on cells undergoing reprogramming to established cell lines and primary cell populations. Both pluripotent stem cell lines — H9 ESCs and established iPSC lines — have similar metabolic and nuclear parameters, as expected. Fibroblasts from human donors (HDFs) had metabolic parameters significantly different from EPCs (**Fig. S2**). This could be because 1) fibroblasts are adherent while starting EPCs are non-adherent, 2) fibroblasts and EPCs have different proliferation rates and energy needs.

Taken together, autofluorescence imaging of NAD(P)H and FAD revealed significant dynamic changes for various cell populations during reprogramming.

### OMI enables the classification of reprogramming cells with high accuracy

To visualize cell states within the entire metabolic and nuclear morphometry dataset, Uniform Manifold Approximation and Projection (UMAP)^65^, a dimension reduction technique, was employed on the multi-dimensional measurements described above. Neighbors were defined through the Jaccard similarity coefficient computed across the metabolic parameters and nuclear parameters. UMAP was chosen over t-distributed stochastic neighbor embedding (t-SNE) since UMAP (**Fig. 2a**) yielded more distinct clusters for two different known cell types — EPCs and iPSCs — than t-SNE (**Fig. S4a**). Moreover, we found that UMAP on our dataset has a higher speed. In addition, UMAP can include non-metric distance functions while preserving the global structure of the data.

**Fig. 2.**
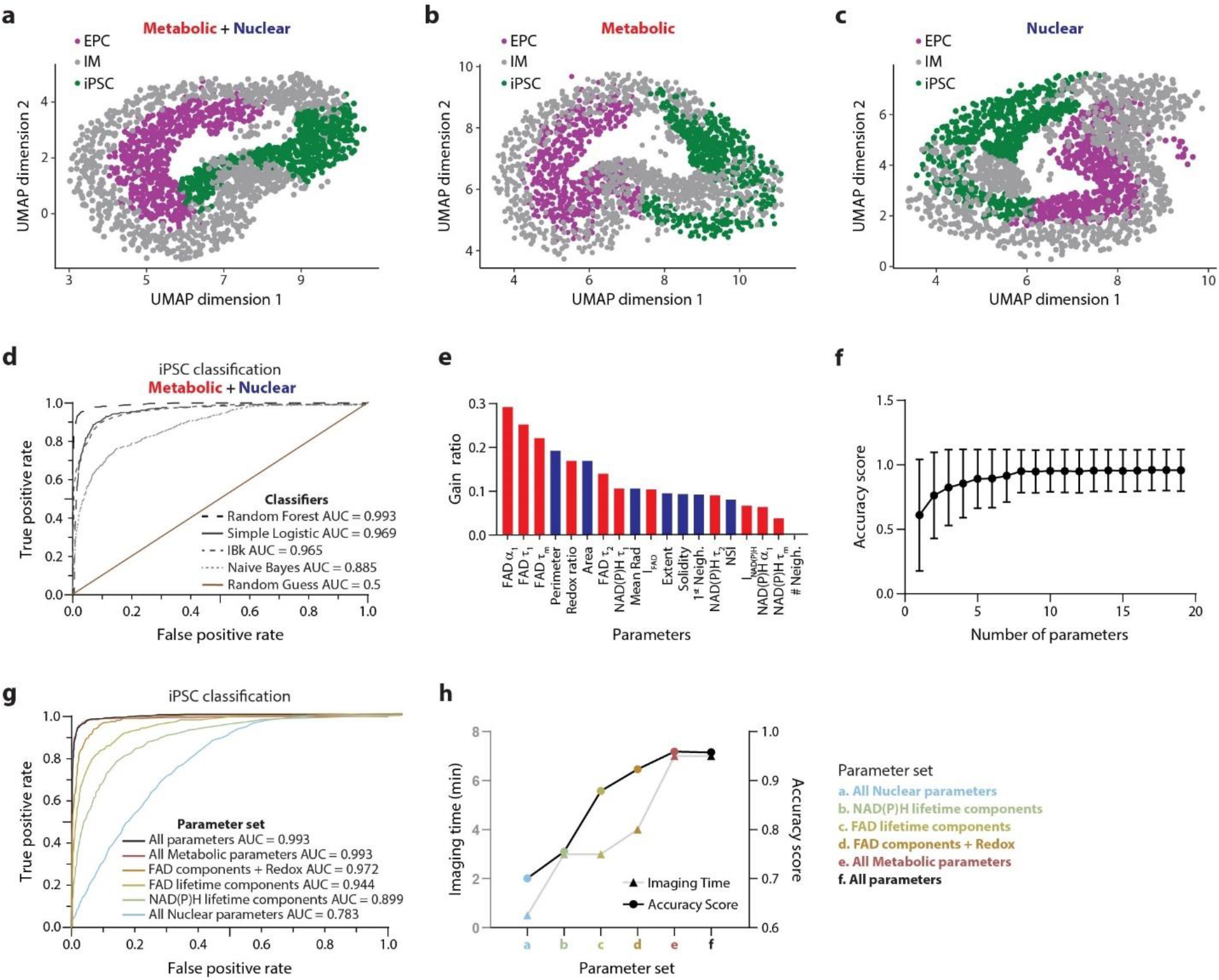
Optical metabolic imaging enables the classification of cells based on their reprogramming status. Uniform Manifold Approximation and Projection (UMAP) dimensionality reduction was performed on **a)** all 11 metabolic and 8 nuclear parameters, **b)** only 11 metabolic parameters, and **c)** only 8 nuclear parameters for each cell, projected onto 2D space and enables separation of different cell types (EPC, IM, and iPSC). Each color corresponds to a different cell type. Data are from three different donors. Each dot represents a single cell, and n =561, 990, and 586 cells for EPC, IM, and iPSC, respectively. **d)** Model performance of the different classifiers (random forest, simple logistic, k-nearest neighbor (IBk), naïve bayes) for iPSCs was evaluated by receiver operating characteristic (ROC) curves using all 11 metabolic and 8 nuclear parameters. The area under the curve (AUC) is provided for each classifier as indicated in the legend. **e)** Parameter weights for random forest classification of EPCs, IMs, and iPSCs using the gain ratio method. Analysis was performed at a single-cell level using three different donors. **f)** Classification accuracy with respect to number of parameters was evaluated based on the gain ratio parameter selection with the random forest model (parameters added from highest to lowest gain ratio in panel **e**. The number of parameters included in the random forest model is indicated on the x-axis. **g)** Model performance of the random forest classifier for iPSCs was evaluated by ROC curves using different metabolic and nuclear parameter combinations as labeled. AUC is provided for each parameter combination as indicated in the legend. **h)** Imaging time (left y-axis) and accuracy score (right y-axis) evaluation of the random forest classifier for different metabolic and nuclear parameter combinations as labeled.

UMAP was next used on subsets of the entire dataset to investigate which measurements were leading to different cell states. Distinct cell populations could be derived from datasets built exclusively from the 11 metabolic parameters (**Fig. 2b**) and datasets built exclusively from the 8 nuclear parameters (**Fig. 2c**). While these UMAP representations revealed some distinct clusters of EPCs, IMs, and iPSCs; UMAP generated using both metabolic and nuclear parameters provided less overlap of cell clusters among EPCs, IMs, and iPSCs (**Fig. 2a**). We also plotted a heatmap representation of the *z-*score of metabolic and nuclear parameters at the donor level (each row is the mean data of a single donor and cell type) to examine heterogeneity arising from individual donors (**Fig. S4b**). Despite donor-to-donor heterogeneity, EPCs and iPSCs could be distinguished based on a combination of 11 metabolic and 8 nuclear parameters.

Next, classification models were developed based on 11 metabolic and 8 nuclear parameters to predict the reprogramming status of cells, i.e., EPCs, IMs, or iPSCs. Supervised machine learning classification (Naïve Bayes, K-nearest neighbor) and regression algorithms (logistic regression, and random forest)^66^ were implemented to test the prediction accuracy for iPSCs when all the metabolic and nuclear parameters are used. To protect against over-fitting, various classification methods were trained using 15-fold cross-validation on single-cell data from three different donors with reprogramming status assigned from morphological characteristics. Further, we tested the various classification methods on data collected from CD71 and Nanog immunofluorescence staining with the same cells from three donors (completely independent and non-overlapping observations). Receiver operator characteristic (ROC; One-vs-Rest) curves of the test data revealed highest classification accuracy for predicting iPSCs (area under the curve, AUC = 0.993), IMs (AUC = 0.993) and EPCs (AUC = 0.999) when a random forest classification model is used (**Fig. 2d, Fig. S4c**,**d**). We thus used the random forest classification model for further analysis in this study.

Gain ratio analysis on the decision tree within this random forest model revealed that FAD lifetime components, FAD α_1_, FAD τ_1,_ and FAD τ_m_, are the most important parameters for classifying the reprogramming status of cells (**Fig. 2e**). This result is consistent with the observation that FAD lifetime components are significantly different among EPCs, IMs, and iPSCs (**Fig. 1, Fig. S2**). We then plotted the accuracy score as a function of the number of parameters (chosen based on the gain ratio values for random forest classifier) used for classification. This plot revealed that the accuracy score increases with the number of parameters until 8 parameters and plateaus thereafter (**Fig. 2f**). Notably, high classification accuracy can be achieved for predicting iPSCs (area under the curve AUC = 0.944), IMs (AUC = 0.968) and EPCs (AUC = 0.987) when using only FAD lifetime variables (FAD τ_m_, τ_1_, τ_2_, α_1_; collected in the FAD channel alone) (**Fig. 2g, Fig. S4e**,**f**). Using only FAD lifetime parameters ensures minimal imaging time of 2.5 min per μFeature (**Fig. 2h**), and no additional reliance on intensity parameters which are associated with higher variability due to the confounding factors of intensity levels (*e*.*g*., throughput due to laser power, detector gain). Hence, FAD lifetime parameters alone are sufficient to predict the reprogramming status of cells.

### Pseudotemporal ordering of single cells resolves cellular transitions

By sampling a process over a time course, single-cell profiles can be used to order cells along a “pseudotemporal” continuum; a method that has helped resolve cellular transitions during development^45,67^. Here we used 11 metabolic and 8 nuclear parameters to construct pseudotime single-cell trajectories of cellular reprogramming using the Monocle3 algorithm^68,69^. Monocle3 is a trajectory inference method that learns combinatorial changes that each cell must go through as a part of a process and subsequently places each cell at its inferred location in the trajectory. The inferred pseudotime trajectories built on our entire dataset consisted of EPCs, IMs, and iPSCs distributed across 10 clusters, 4 branching events, and a disconnected branch (**Fig. 3a-c**). The primary trajectory colored by pseudotime and actual reprogramming time points showed ordering from EPCs to IMs to iPSCs as expected (**Fig. 3b, Fig. S5a**).

**Fig. 3.**
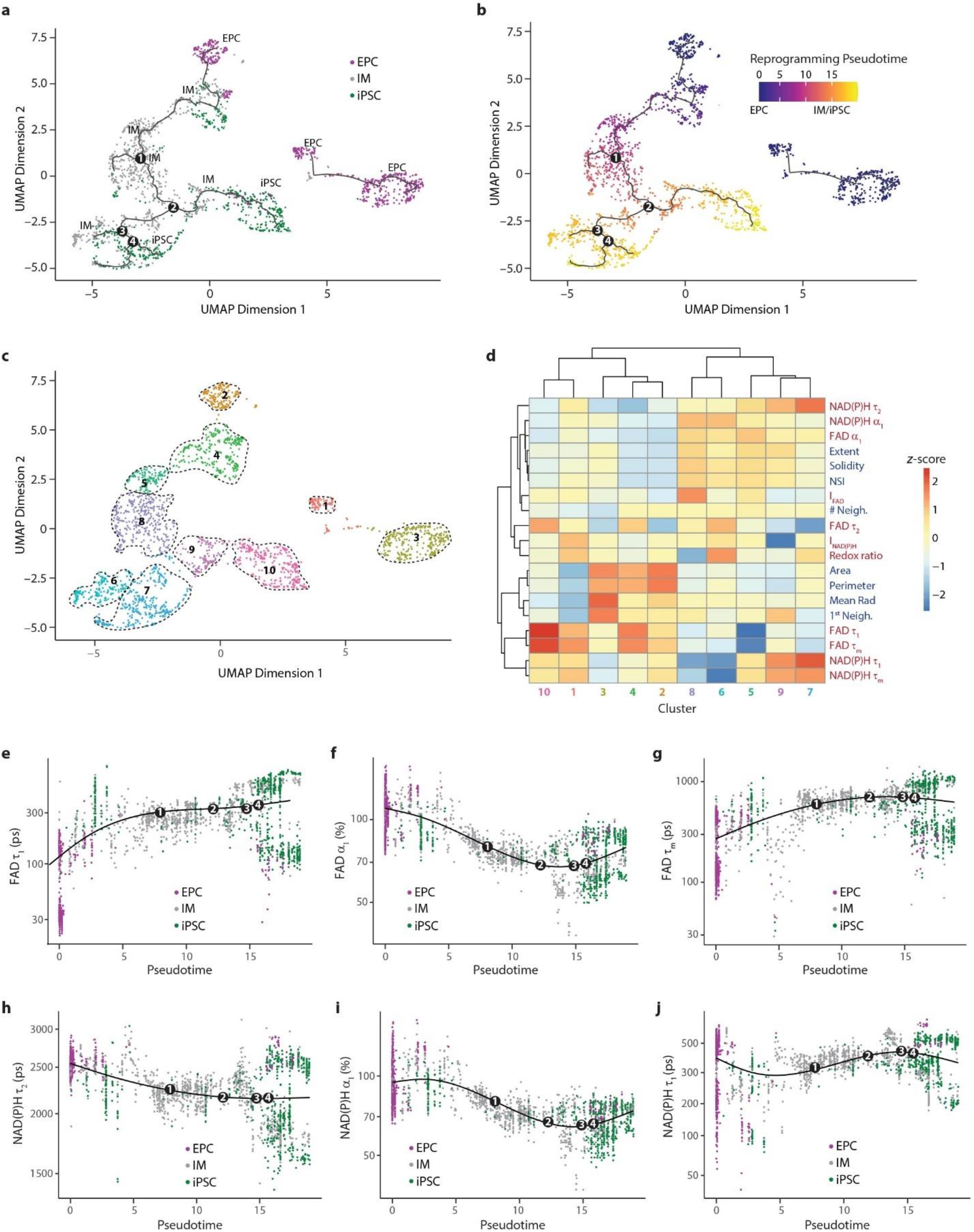
Inference of single-cell reprogramming trajectories reveals heterogeneity during reprogramming. Trajectory analysis of reprogramming EPCs constructed from the metabolic and nuclear parameters based on UMAP dimension reduction using Monocle3 revealed four branch points, colored by **a)** cell type and **b)** pseudotime. **c)** Monocle UMAP plots showing clustering of reprogramming EPCs. Samples were grouped into 10 clusters. Cells colored by cluster. **d)** Heatmap representing the metabolic and nuclear parameters of 10 clusters. Each column is a separate cell group based on the generated clusters and each row represents a single metabolic or nuclear parameter. *Z*-score = (*μ*_*observed*_−*μ*_*row*_)/σ_*row*_, where μ_observed_ is the mean value of each parameter for each cell; μ_row_ is the mean value of each parameter for all cells together, and σ_row_ is the standard deviation of each parameter across all cells. Dot plots indicating the expression of **e)** FAD τ_1_, **f)** FAD α_1_, **g)** FAD τ_m_, **h)** NAD(P)H τ_2_, **i)** NAD(P)H α_1_ and, **j)** NAD(P)H τ_2_ along the pseudotime. Smooth lines are composed of multiple dots representing the mean expression level at each pseudotime, regardless of the cell type. Four branch points are labeled on the smooth lines.

Trajectory inference indicated that the starting EPCs were heterogeneous and occupied three clusters (**Fig. 3c**; clusters: 1, 2, 3). While cluster 2 consists of starting EPCs that undergo reprogramming, clusters 1 and 3 constituted the disconnected branch with EPCs that were not permissive to reprogramming. iPSCs predominantly occupied two clusters (clusters: 7, 10) irrespective of the reprogramming timepoint, while IMs belonged to several clusters (clusters: 4, 5, 6, 8, 9) with clusters 6 and 8 concentrated at the unsuccessful reprogramming branches (**Fig. 3c**). Overall, these various trajectories provide a detailed map of several cases of reprogramming heterogeneity within human cells. For example, cells that advance right at branch points 1, 2, and 3 (**Fig. 3a,b**) completely reprogram to iPSCs within 25 days of reprogramming initiation while cells that proceed left at branch points 1 and 3 (**Fig. 3a,b**) remain at the intermediate stage.

Subsequent heatmap analysis (**Fig. 3d**) on the clusters in the single-cell reprogramming trajectory map revealed that the clusters exhibited correlation patterns based on their reprogramming status i.e., EPCs (clusters: 2) have a high correlation to early IMs (cluster: 4), while late IMs (clusters: 5,6,8,9) demonstrate high correlation to iPSCs (clusters: 7) (**Fig. 3d**). When we compared IMs that undergo reprogramming (cluster: 9) and the IMs that do not reprogram to iPSCs (cluster: 6), we noted differences in their NAD(P)H lifetime components, indicating that these parameters might play a role in determining reprogramming cell fate. To further examine the parameters that distinguished the cell clusters, we performed spatial correlation analysis using Moran’s I^70^, which is a statistic that reports whether cells at nearby positions on a trajectory will have similar (or dissimilar) expression levels for a given parameter (**Fig. S5b**). When the parameters were ranked by Moran’s I, FAD lifetime parameters (FAD τ_1_, τ_2_, τ_m_) were most important in distinguishing clusters followed by NAD(P)H lifetime parameters [NAD(P)H τ_2_, α_1,_ τ_1_], in agreement with expression level maps (**Fig. S5c-h**). This result is consistent with high gain ratio values for FAD lifetime parameters (**Fig. 2e**) and the observation FAD lifetime parameters are significantly different among EPCs, IMs, and iPSCs (**Fig. 1d**).

While FAD parameters are important in distinguishing EPCs, IMs, and iPSCs, NAD(P)H parameters are key for determining the eventual reprogramming fate of cells. When we plotted the identified important metabolic parameters as a function of pseudotime, we observed that they undergo biphasic changes during reprogramming that could be representative of the OXPHOS burst (**Fig. 3e-j**). These pseudotime trajectories complement the UMAP visualizations (**Fig. 2a-c)** by providing higher temporal resolution of changes occurring during reprogramming.

### Isolation of high-quality iPSCs

While visualizing reprogramming heterogeneity at a high temporal resolution and single-cell resolution with our methods can be insightful, the terminal goal of any reprogramming platform is to successfully isolate iPSCs that can be used for downstream applications. As proof-of-concept, we used a combination of OMI, μCP platform, and machine learning models developed in this study to isolate high-quality iPSCs (**Fig. 4**). First, we tracked the metabolic and nuclear parameters of μFeatures throughout the reprogramming time course using OMI (**Fig. 4a**). Second, we employed our random forest classification model to predict the reprogramming status of the tracked μFeatures (**Fig. 4b**). Third, we inferred the pseudotimes during the reprogramming time course to monitor the progress of the μFeatures along the reprogramming trajectory (**Fig. 4c**). Finally, we performed immunostaining on the μFeatures, which showed that the reprogramming status predictions made by the machine learning models correlated well with the actual staining (**Fig. 4d**).

**Fig. 4.**
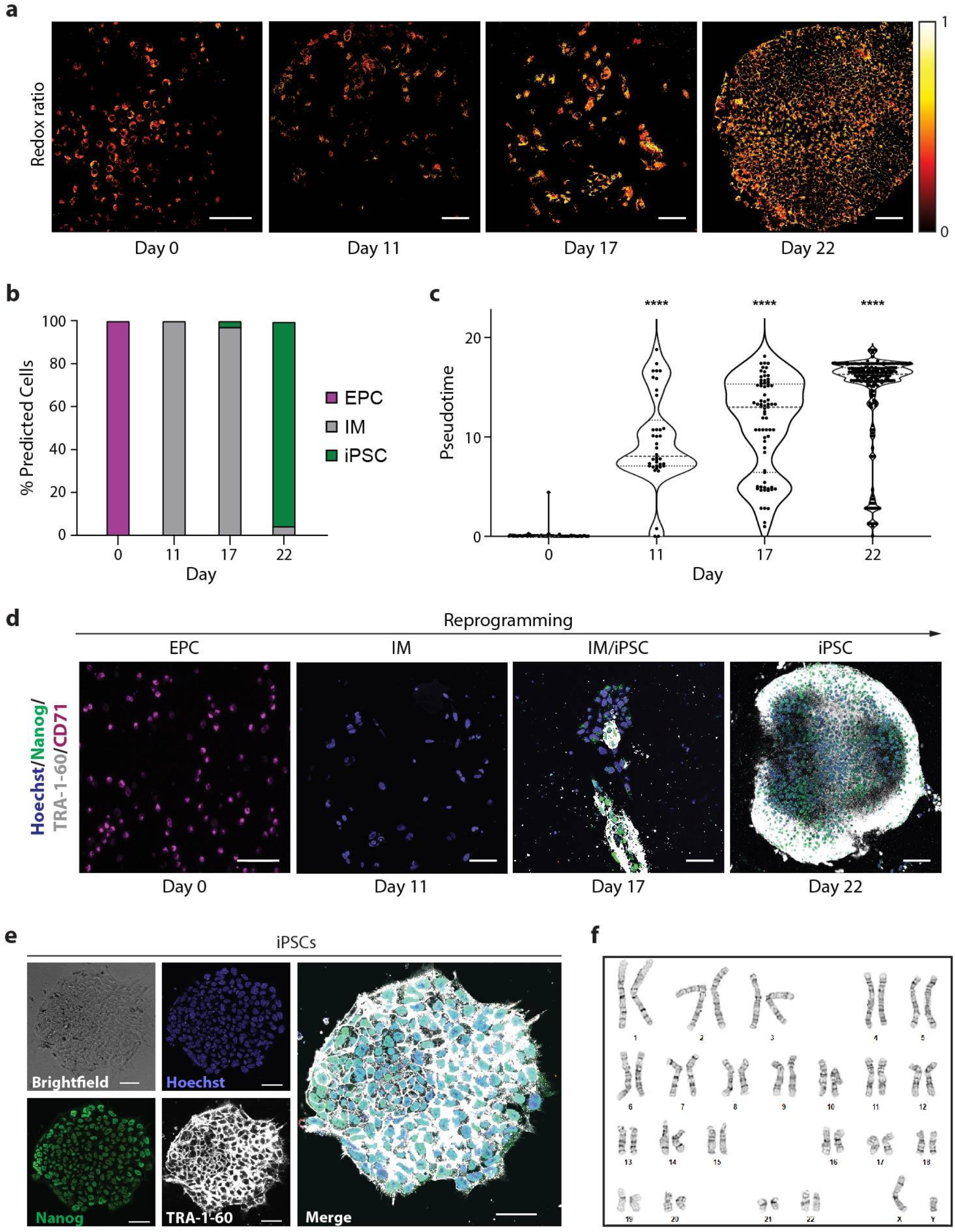
Optical metabolic imaging of μFeatures aids in the identification and isolation of iPSC populations. **a)** Representative optical redox ratio images of a single μFeature at different days through the reprogramming time course. Color bar is indicated on the right. Scale bar, 100 μm. **b)** Stacked column bar graph showing the variation in distribution of cell types during reprogramming as predicted by random forest classifier using all metabolic and nuclear parameters. The color of the bar corresponds to the cell type and the height of the bar represents the percentage of cell types. **c)** Violin plots showing the distribution of reprogramming pseudotime of single cells within a μFeature as a function of the actual reprogramming timepoint. Dashed lines indicate median and dotted lines indicate the interquartile range. Statistical significance was determined by one-way analysis of variance (ANOVA) using the Kruskal-Wallis test for multiple comparisons; ns = p ≥0.05, * for p <0.05, ** for p <0.01, *** for p <0.001, **** for p <0.0001). **d)** Representative images of cell subpopulations on μFeatures at different days through the reprogramming time course, stained using antibodies against Hoechst (blue), TRA-1-60 (white), Nanog (green), and CD71 (magenta). Scale bar, 100 μm. **e)** Representative image of iPSC colony isolated from a μFeature, stained with Hoechst (nuclear dye), TRA-1-60, and Nanog (pluripotency markers). Scale bar, 50 μm. **f)** iPSCs derived from μCP substrates show normal karyotype suggesting that no major chromosome abnormality was present within the cells after reprogramming.

We then isolated iPSCs from the μCP culture platform based on the predictions made by the random forest classification model. The physical separation of micropatterns from one another, combined with a high fraction of predicted iPSCs, even up to 100% throughout the μFeature, resulted in easy picking and isolation of completely reprogrammed iPSCs. We further confirmed that the isolated iPSCs expressed pluripotency markers (**Fig. 4e**) and showed no genomic abnormalities (**Fig. 4f**), indicating that our reprogramming platform can be used to generate genetically-stable iPSC lines.

## Discussion

Here, we report a non-invasive, high-throughput, quantitative, and label-free imaging platform to predict the reprogramming outcome of EPCs by combining micropatterning, live-cell autofluorescence imaging, and automated machine learning. We can predict the reprogramming status of EPCs at any timepoint during reprogramming with a prediction accuracy of ∼95% and model performance of ∼0.99 (AUC of ROC) using a random forest classification model with 11 metabolic parameters and 8 nuclear parameters (**Fig. 2g, Fig. S4c-f**). Additionally, we provide a single cell roadmap of EPC reprogramming, which reveals diverse cell fate trajectories of individual reprogramming cells (**Fig. 3**).

Recent evidence indicates that metabolic changes during reprogramming include decreasing OXPHOS and increasing glycolysis^26,71^, along with a transient hyper-energetic metabolic state, called OXPHOS burst. This OXPHOS burst occurs at an early stage of reprogramming and shows characteristics of both high OXPHOS and high glycolysis, which could be a regulatory cue for the overall shift of reprogramming^28,29,72,73^. These changes are accompanied by alterations in the amounts of corresponding metabolites and have been confirmed by genome-wide analyses of gene expression, protein levels, and metabolomic profiling^74–77^. The shifts in cellular metabolism affect enzymes that control epigenetic configuration^78^, which can impact chromatin reorganization and provide a basis for changes in nuclear morphology as well as gene expression during reprogramming ^34,79–82^. Consistent with these studies, the redox ratio increases during reprogramming (**Fig. 1d**), which could be indicative of increased glycolysis during reprogramming^83^.

The changes in NAD(P)H and FAD lifetime parameters that occur during reprogramming (**Fig. 1, Fig. S2**) could reflect changes in quencher concentrations, such as oxygen, tyrosine, or tryptophan, or changes in local temperature and pH^35,84,85^. Specifically, the biphasic changes in the metabolic and nuclear parameters could due to the increased production of ROS by mitochondria^35,58,86,87^ during the OXPHOS burst. The generated ROS further serves as a signal to activate Nuclear Factor (erythroid derived 2)-like-2 (NRF-2), which then induces hypoxia-inducible factors (HIFs) that promote glycolysis during reprogramming by increasing the expression levels of the glycolysis-related genes^25,73,76^.

Moreover, the importance of FAD parameters for distinguishing various reprogramming cell types (**Fig. 1d, Fig. 2e**) could point to the significant changes in the mitochondrial environment during reprogramming. The differences in NAD(P)H lifetime parameters among IMs that successfully undergo reprogramming and the ones that do not (**Fig. 3d**), may suggest the role for NAD(P)H in impacting reprogramming barriers.

The classification analysis revealed that models trained on all 11 metabolic and 8 nuclear parameters yielded the highest accuracy for the classification of reprogramming status of cells. Random forest classification using only FAD lifetime parameters yielded comparatively high ROC AUC values (**Fig. 2g, Fig. S4e**,**f**). Additionally, FAD lifetime parameters were more accurate for predicting reprogramming status than using nuclear parameters alone, which can be obtained using widefield or confocal fluorescence microscopy. Imaging only FAD lifetime parameters instead of imaging all the parameters significantly reduced the time of imaging from 7 min to 2.5 min per μFeature (**Fig. 2h**). This is especially helpful when assessing multiple μFeatures for iPSC quality at a manufacturing scale, and lifetime measurements benefit from fewer confounding factors and less variability compared with intensity measurements.

Our single-cell reprogramming trajectory maps based on metabolic and nuclear parameters (**Fig. 3**) could indicate that the reprogramming process proceeds by a combination of elite and stochastic models^88^. While a fraction of starting EPCs are refractory towards reprogramming supporting the elite model of reprogramming^89^, there is also a fraction of intermediate cells at various stages of reprogramming that do not completely reprogram to iPSCs corroborating the stochastic model of reprogramming^90^.

Much of the current work to understand the heterogeneity during reprogramming relies on bulk analysis^91–94^ or single-cell analysis^95–103^ techniques. While bulk samples obscure variability in both the starting cell population and during fate conversion, owing to the variable kinetics and low efficiency of reprogramming; single-cell techniques disrupt the cells’ microenvironment, resulting in significant changes in the biophysical properties of cells undergoing reprogramming. Our methods overcome these challenges with the combination of a μCP culture platform, OMI, and machine learning. Firstly, the μCP platform ensures an intact microenvironment for reprogramming cells while enabling single-cell analysis. Secondly, OMI provides single-cell measurements non-destructively to assess the influence of neighboring cells and provides high temporal resolution for time-course studies of reprogramming. Finally, machine learning with trajectory inference is applied here to a new type of cellular measurement, single cell metabolism. These methods excel in analyzing time course data containing asynchronous processes within cells — as seen in prior studies with flow cytometry and gene expression data^68,75,101,102,104^. Machine learning here overcomes the problems of reprogramming trajectories built based on absolute time points that disregard the asynchrony of the reprogramming process. Overall, these methods could aid in the identification of somatic cells or early reprogramming cells that are refractory towards reprogramming and thus increase the success rate of iPSC generation from patient-derived primary cells or cell lines.

These methods could be adapted for industrial-scale, GMP-compliant manufacturing system. First, the μCP platform involves direct ECM printing onto optically clear substrates (**Fig. S1a**) and does not involve any gold coating, unlike traditional microcontact printing methods^105,106^. Therefore, the μCP platform is cost-effective and relatively simple because it does not require cleanroom access. Second, we isolated fully pluripotent iPSCs without any genomic abnormalities (**Fig. 4f**) using this μCP platform. The use of EPCs with episomal reprogramming plasmids are likely to generate genetically-stable iPSCs devoid of reprogramming factors, as based on prior studies with EPCs^107,108^ and episomal plasmids^52,109,110,111^. Plus, our μCP platform is xeno-free and feeder-free to eliminate the reprogramming inconsistencies arising from the undefined nature of xeno-components of other reprogramming culture systems. Third, the autofluorescence imaging technique is label-free, unlike other common methods to study metabolism such as electron microscopy, immunocytochemistry, and colorimetric metabolic assays. Autofluorescence imaging provides non-destructive real-time monitoring of live cells with lower sample phototoxicity compared to single-photon excitation^112^. Taken together, the processes of μCP platform fabrication, reprogramming, cell culture, autofluorescence imaging, iPSC identification based on machine learning models and iPSC isolation can all be automated and extended to different reprogramming methods^113^ (*e*.*g*., mRNA, Sendai virus), to other starting cell types (*e*.*g*., fibroblasts, keratinocytes), to other parameters (*e*.*g*., cell morphology^19–21^, mitochondrial structure^71,77,114^), and to other processes (e.g., differentiation^115–118^).

Some of the limitations of our current approach include two-dimensional imaging, culture duration, and per-μFeature image analysis. First, comprehensive three-dimensional imaging of each μFeature could provide maps at higher resolution to further dissect the metabolic and nuclear changes occurring throughout the entire depth of the reprogramming cultures. Second, there is a limited duration of culture before cells overgrow within the μFeature. This could also result in cell detachment from the μFeature, which is difficult to image with OMI. For the reprogramming experiments described here, circular features with 300 μm radius have been used for ∼25 days of culture, although the cell seeding density or micropatterned geometry^34,43,105^ could be easily changed. Finally, imaging analysis was performed at different reprogramming timepoints on a per-μFeature basis. However, tracking single cells within the μFeature during reprogramming using cell tracking algorithms^119^ could provide deeper insights into metabolic and nuclear changes during reprogramming.

Overall, we developed a high-throughput, non-invasive, rapid, and quantitative method to predict the reprogramming status of cells and study reprogramming heterogeneity. Our studies indicate that OMI can predict the reprogramming status of cells, which could enable real-time monitoring during iPSC manufacturing, thereby aiding in the identification of high-quality iPSCs in a timely and cost-effective manner. Similar technologies could impact other areas of cell manufacturing such as direct reprogramming, differentiation^115^, and cell line development.

## Materials and Methods

### EPC isolation and cell culture

EPCs were isolated from fresh peripheral human blood that was obtained from healthy donors (Interstate Blood Bank, Memphis, TN). Blood was processed within 24 hours of collection, where hematopoietic progenitor cells were extracted from whole blood using negative selection (RosetteSep; STEMCELL Technologies) and cultured in polystyrene tissue culture plates in erythroid expansion medium (STEMCELL Technologies) for 10 days to enrich for EPCs. Enriched EPCs from Day 10 were examined by staining with APC Anti-Human CD71 antibody (334107; Biolegend; 1:100) and incubating for 1 hour at room temperature. Data were collected on Attune Nxt flow cytometer and analyzed with FlowJo.

### Micropattern Design and PDMS stamp production

First, a template with the feature designs was created in AutoCAD (Autodesk). The template was then sent to the Advance Reproductions Corporation, MA for the fabrication of a photomask, and a 6-inch patterned Si wafer was fabricated by the Microtechnology Core, University of Wisconsin-Madison, WI^120^. Using soft photolithography techniques, the Si wafer was spin-coated with a SU-8 negative photoresist (MICRO CHEM) and exposed to UV light. The Si mold was then developed for 45 minutes in SU-8 developer (Sigma) which yielded features with a height of 150 μm. The Si mold was then washed with acetone and isopropyl alcohol.

Elastomeric stamps used for microcontact printing were generated by standard soft lithographic techniques. The silicon mold was rendered inert by overnight exposure in vapors of (tridecafluoro-1, 1, 2, 2-tetrahydrooctyl) trichlorosilane. Poly-dimethylsiloxane (Sylgard 184 silicone elastomer base, 3097366-1004, Dow Corning; PDMS) was prepared at a ratio of 1:10 curing agent (Sylgard 184 silicone elastomer curing agent, 3097358-1004, Dow Corning) and degassed in a vacuum for 30 minutes. The PDMS was then poured over the SU-8 silicon mold on a hot plate and baked at 60°C overnight to create the PDMS stamp.

### μCP Well Plate construction

Microcontact patterned (μCP) substrates were constructed based on previous studies^47,48,121^. In brief, polydimethylsiloxane (PDMS) stamps with 300 μm radius circular features were coated with Matrigel (WiCell Research Institute) for 24 h. After 24 h, the Matrigel-coated PDMS stamp was dried with N_2_ and placed onto 35 mm cell culture treated ibiTreat dishes (81156; Ibidi). A 50 g weight was added on top of the PDMS stamps to ensure even pattern transfer from the Matrigel-coated PDMS stamp to the ibiTreat dish. This setup was incubated for 2 h at 37°C. The 35 mm ibiTreat dish was then backfilled with PLL (20 kDa)-*g*-(3.5)-PEG (2 kDa) (Susos), a graft polymer solution in with a 20 kDa PLL backbone with 2 kDa PEG side chains, and a grafting ratio of 3.5 (mean PLL monomer units per PEG side chain), by using 0.1 mg/mL solution in 10 mM HEPES buffer for 30 min at RT. The ibiTreat dish was then washed with PBS and exposed to UV light for 15 min for sterilization to yield the micropatterned substrate.

### Reprogramming

Day 10 EPCs were electroporated with four episomal reprogramming plasmids encoding Oct4, shRNA knockdown of p53 (#27077; Addgene); Sox2, Klf4 (#27078; Addgene); L-Myc, Lin28 (#27080; Addgene); miR302-367 cluster (#98748; Addgene), using the P3 Primary Cell 4D-Nucleofector Kit (Lonza) and the EO-100 program^51,52^. Electroporated EPCs were seeded onto micropatterned substrates with erythroid expansion medium (STEMCELL Technologies) at a seeding density of 2000k cells/dish. Cells were supplemented with ReproTeSR (STEMCELL Technologies) on alternate days starting from Day 3 without removing any medium from the well. On Day 9, the medium was entirely switched to ReproTeSR, and the ReproTeSR medium was changed daily starting from Day 10.

### Isolation of iPSCs

To isolate high-quality iPSC lines, candidate colonies were picked from micropatterns using a 200 μL micropipette tip and transferred to Matrigel-coated polystyrene tissue culture plates in mTeSR1 media (WiCell Research Institute). If additional purification was required, one additional manual picking step with a 200 μL micropipette tip was performed. During picking and subsequent passaging, the culture media was often supplemented with the Rho kinase inhibitor Y-27632 (Sigma-Aldrich) at a 10 μM concentration to encourage cell survival and establish clonal lines. iPSCs obtained from EPCs were maintained in mTeSR1 media on Matrigel-coated polystyrene tissue culture plates and passaged with ReLeSR (STEMCELL Technologies) every 3-5 days. All cells were maintained at 37ºC and 5% CO_2_.

### Antibodies and Staining

All cells were fixed for 15 minutes with 4% paraformaldehyde in PBS (Sigma-Aldrich) and permeabilized with 0.5% Triton-X (Sigma-Aldrich) for >4 hours at room temperature before staining. Hoechst (H1399; Thermo Fisher Scientific, Waltham, MA) was used at 5 μg/mL with 15 min incubation at room temperature to stain nuclei. Primary antibodies were applied overnight at 4°C in a blocking buffer of 5% donkey serum (Sigma-Aldrich) at the following concentrations: Anti-Laminin (L9393; Sigma-Alrich) 1:500; TRA-1-60 (MAB4360; EMD Millipore, Burlington, MA) 1:100; Nanog (AF1997; R&D Systems) 1:200; CD71 (334107; Biolegend) 1:100. Secondary antibodies were obtained from Thermo Fisher Scientific and applied in a blocking buffer of 5% donkey serum for one hour at room temperature at concentrations of 1:400 – 1:800. A Nikon Eclipse Ti epifluorescence microscope was used to acquire single 10x images of each micropattern, and a Nikon AR1 confocal microscope was used to acquire 60x stitched images of each micropattern using the z-plane closest to the micropatterned substrate for reprogramming studies.

### Autofluorescence imaging of NAD(P)H and FAD

Fluorescence lifetime imaging (FLIM) was performed at different time points during reprogramming by an Ultima two-photon microscope (Bruker) composed of an ultrafast tunable excitation laser source (Insight DS+, Spectra-Physics) coupled to a Nikon Ti-E inverted microscope with time-correlated single-photon counting electronics (SPC-150, Becker & Hickl). The laser source enables sequential excitation of NAD(P)H at 750 nm and FAD at 890 nm. NAD(P)H and FAD images were acquired through 440/80 nm and 550/100 nm bandpass filters (Chroma), respectively, using Gallium arsenide phosphide (GaAsP) photomultiplier tubes (PMTs; H7422, Hamamatsu). The laser power at the sample was approximately 3.5 mW for NAD(P)H and 6 mW for FAD. Lifetime imaging using time-correlated single-photon counting electronics (SPC-150, Becker & Hickl) was performed within Prairie View Atlas Mosaic Imaging (Bruker Fluorescence Microscopy) to capture the entire μFeature. Fluorescence lifetime decays with 512-time bins were acquired across 512 × 512-pixel images with a pixel dwell time of 4.8 μs and an integration period of 60 seconds. Photon count rates were ∼1-5 × 10^5^ and monitored during image acquisition to ensure that no photobleaching occurred. All samples were placed on a stage-op incubator and illuminated through a 40×/1.15 NA objective (Nikon). The short lifetime of red-blood-cell fluorescence at 890 nm was used as the instrument response function and had a full-width half maximum of 240 ps. A YG fluorescent bead (τ = 2.13 ± 0.03 ns, n= 6) was imaged daily as a fluorescence lifetime standard^35,122^.

### Image analysis

Fluorescence lifetime decays were analyzed to extract fluorescence lifetime components via SPCImage software (Becker & Hickl). A threshold was used to exclude pixels with low fluorescence signals (that is, background). A bin of 3×3 pixels was used to maintain spatial resolution, the fluorescence lifetime decay curve was convolved with the instrument response function and fit to a two-component exponential decay model, I(t) = α_1_e^-t/τ^_1_ + α_2_e^-t/τ^_2_+ C, where *I*(*t*) is the fluorescence intensity as a function of time *t* after the laser pulse, *α*_1_ and *α*_2_ are the fractional contributions of the short and long lifetime components, respectively (that is, *α*_1_ + *α*_2_ = 1), *τ*_1_ and *τ*_2_ are the short and long lifetime components, respectively, and *C* accounts for background light. Both NAD(P)H and FAD can exist in quenched (short lifetime) and unquenched (long lifetime) configurations^39,40^; the fluorescence decays of NAD(P)H and FAD are therefore fit to two components. Fluorescence intensity images were generated by integrating photon counts over the per-pixel fluorescence decays.

### Images were analyzed at the single-cell level to evaluate cellular heterogeneity^123^

A pixel classifier was trained on 15 images using ilastik^59^ software to identify the pixels within the nuclei in NAD(P)H images. An object classifier was then used to identify the nuclei in NAD(P)H images using the pixel classifier along with the following parameters: Method = Simple, Threshold = 0.3, Smooth = 1, Size Filter Min = 15 pixels, Size Filter Max = 500 pixels. A customized CellProfiler^60^ pipeline was then used to obtain metabolic and nuclear parameters. The CellProfiler pipeline applied the following steps: Primary objects (nuclei) were inputted from ilastik. Secondary objects (cells) were then identified in the NAD(P)H intensity image by outward propagation of the primary objects. Cytoplasm masks were determined by subtracting the nucleus mask from the cell mask. Cytoplasm masks were applied to all images to determine single-cell redox ratio and NAD(P)H and FAD lifetime parameters. A total of 11 metabolic parameters were analyzed for each cell cytoplasm (**Fig. S1d**): NAD(P)H intensity (Inad(p)h), NAD(P)H α_1_, NAD(P)H τ_1_, NAD(P)H τ_2_, NAD(P)H mean lifetime (τ_m_ = α_1_τ_1_ + α_2_τ_2_), FAD intensity (Ifad), FAD α_1_, FAD τ_1_, FAD τ_2_, FAD τ_m_, optical redox ratio [Inad(p)h / (Inad(p)h + Ifad)]. A total of 8 nuclear parameters were analyzed for each nucleus: area, perimeter, mean radius (MeanRad), nuclear shape index (NSI), solidity, extent, number of neighbors (#Neigh), distance to closest neighbor (1stNeigh).

Representative images of the optical redox ratio, NAD(P)H *τ*_m_ and FAD *τ*_m_ were computed using the Fiji software.

### UMAP Clustering

Clustering of cells across EPCs, IMs, and iPSCs was represented using Uniform Manifold Approximation and Projection (UMAP). UMAP dimensionality reduction^65^ was implemented using R on all 11 OMI parameters (optical redox ratio, NAD(P)H τ_m_, τ_1_, τ_2_, α_1_, α_2_; FAD τ_m_, τ_1_, τ_2_, α_1_, α_2_) and/or all 8 nuclear parameters (Area, Perimeter, MeanRad, NSI, Solidity, Extent, #Neigh, 1stNeigh) for projection in 2D space. The following parameters were used for UMAP visualizations: “n _neighbors”: 20; “min_dist”: 0.3, “metric”: Jaccard, “n_components”: 2.

### Z-score hierarchical clustering

Z-score of each metabolic and nuclear parameter for each cell was calculated. Z-score = (*μ*_*observed*_−*μ*_*row*_)/σ_*row*_, where μ_observed_ is the mean value of each parameter for each cell; μ_row_ is the mean value of each parameter for all cells together, and σ_row_ is the standard deviation of each parameter across all cells. Heatmaps of z-scores for all OMI variables were generated to visualize differences in each parameter between different cells. Dendrograms show clustering based on the similarity of average Euclidean distances across all variable z-scores. Heatmaps and associated dendrograms were generated in Python.

### Classification methods

Random forest, Simple Logistic, k-nearest neighbor (IBk), and naïve bayes classification methods were trained to classify reprogramming cells into EPCs, IMs, and iPSCs using Weka software^124^. All data were randomly partitioned into training and test datasets using 15-fold cross-validation for training and test proportions of 93.3% (1994 cells) and 6.7% (143 cells), respectively. Each model was replicated 100 times; new training and test data were generated before each iteration. Parameter weights for metabolic and nuclear parameters were extracted using the GainRatioAttributeEval function in Weka to determine the contribution of each variable to the trained classification models. One-vs-Rest receiver operating characteristic (ROC) curves were generated to evaluate the classification model performance on the classification of test set data and are the average of 100 iterations of data that was randomly selected from training and test sets. All of the ROC curves displayed were constructed from the test datasets using the model generated from the training data sets.

### Karyotyping

Cells cultured for at least 5 passages were grown to 60-80% confluence and shipped for karyotype analysis to WiCell Research Institute, Madison, WI. G-banded karyotyping was performed using standard cytogenetic protocols^125^. Metaphase preparations were digitally captured with Applied Spectral Imaging software and hardware. For each cell line, 20 GTL-banded metaphases were counted, of which a minimum of 5 was analyzed and karyotyped. Results were reported in accordance with guidelines established by the International System for Cytogenetic Nomenclature 2016^126^.

### Statistics

p-values were calculated using the non-parametric Kruskal-Wallis test for multiple unmatched comparisons with GraphPad Prism software. Statistical tests were deemed significant at α≤0.05. Technical replicates are defined as distinct μFeatures within an experiment. Biological replicates are experiments performed with different donors. No a priori power calculations were performed.

## Acknowledgments

We thank members of the Saha lab for helpful discussion and comments on the manuscript. Further, we thank members of the Skala lab for discussions about machine learning classification methods. We also thank plasmid depositors to Addgene, and James Thomson’s lab for sharing the miR302-367 plasmid. We thank the Interstate Blood Bank for supplying fresh blood, Jose Jimenez for manufacturing the silicon mold, and Brett Napiwocki for help with PDMS stamp preparation. We thank Matthew Stefely for the assistance with figure illustrations. We acknowledge generous financial support from the National Science Foundation (CBET-1350178), National Institute for Health (U01 EY032333, R01 CA211082, R01 CA205101, R01 CA185747), University of Wisconsin-Madison Stem Cell and Regenerative Medicine Center, Wisconsin Alumni Research Foundation, Wisconsin Institute for Discovery, and the Morgridge institute for Research.

## Author contributions

The project was conceptualized by K.M. and K.S. The experiments were designed by K.M. K.S., T.M.H and M.C.S., and were carried out by K.M., T.M.H. and J.R. Data analysis was performed by K.M., G.A.B. and E.C.G. The manuscript was written and edited by K.M., K.S. and M.C.S.

## Data and materials availability

All data needed to evaluate conclusions in the paper are included in the paper and/or Supplementary Materials. Additional data related to this paper may be requested from the authors.

