## Supplementary Material for "Label-free imaging to track reprogramming of human somatic cells"

**This PDF file includes:**

Figs. S1 to S5

**Other Supplementary Materials for this manuscript include the following:**

Data S1 to S2

Code S1

### Supplementary Figures

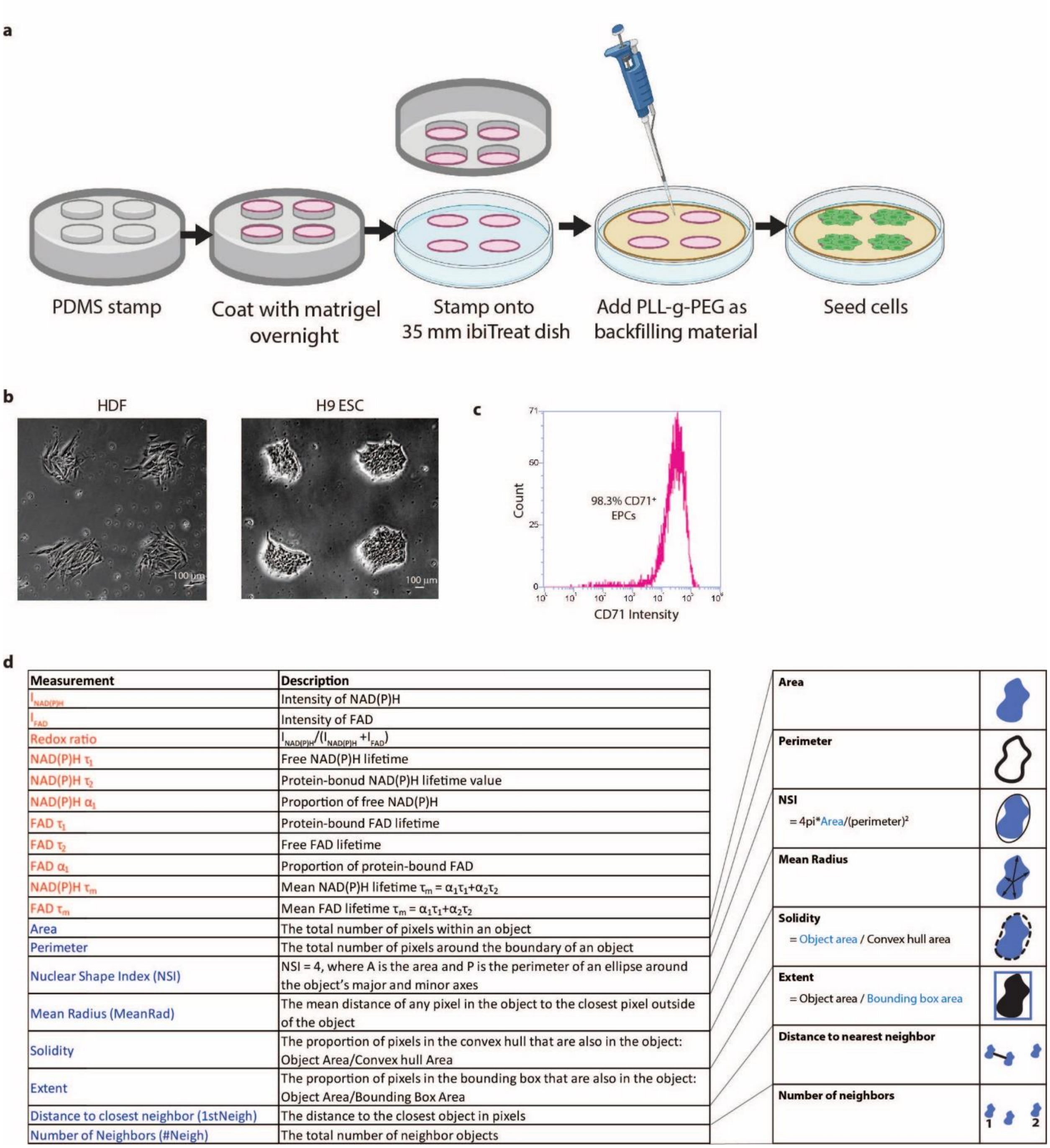


**Fig. S1. Micropatterned substrates enable controlled cell adhesion. a)** Schematic showing the fabrication of µCP substrates. PDMS mold is coated with Matrigel and stamped onto a 35 mm ibiTreat dish. PLL-g-PEG solution is then added to backfill the non-printed regions. Fabricated micropatterned substrates are then ready to be seeded with cells. **b)** Representative images of HDFs and H9 ESCs adhered to 300 µm radius circular µFeatures on micropatterned substrates. Scale bar, 100 µm. **c)** Flow cytometry histogram indicating the percentage of $CD71^+$ cells after 10 days of EPC culture and before electroporation with reprogramming plasmids. **d)** Description of the 11 metabolic and 8 nuclear parameters that were measured. These parameters were obtained after processing NAD(P)H and FAD images using an image analysis pipeline described in Figure 1b.

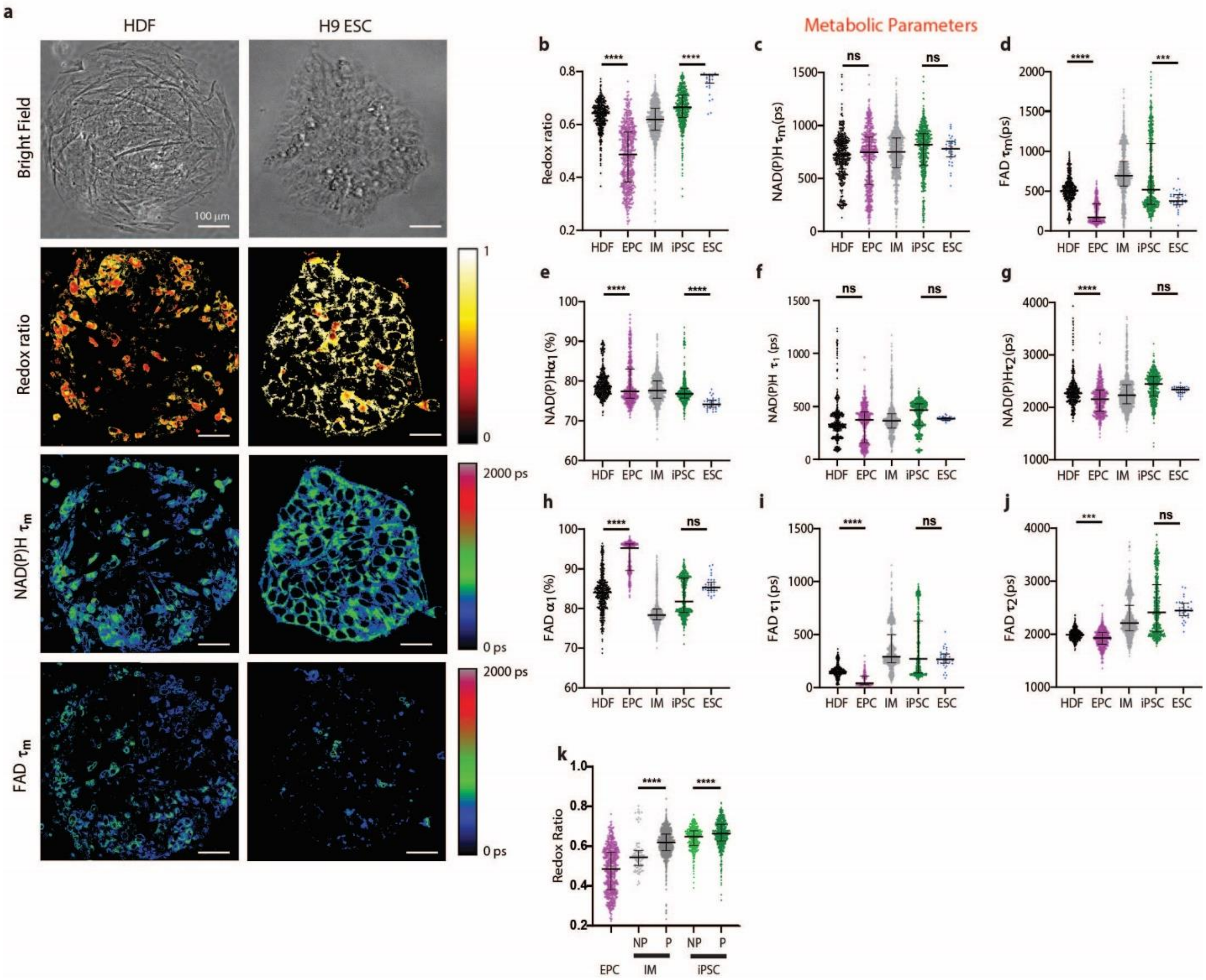


**Fig. S2. Metabolic parameter changes during reprogramming of EPCs. a)** Representative optical redox ratio, NAD(P)H $\tau_m$ and FAD $\tau_m$ images for HDFs and H9 ESC. Color bars are indicated on the right. Scale bar, 100 µm. Single-cell quantitative analysis of **b)** optical redox ratio [$I_{NAD(P)H}/(I_{FAD}+I_{NAD(P)H})$], **c)** NAD(P)H $\tau_m$, **d)** FAD $\tau_m$, **e)** NAD(P)H $\alpha_1$, **f)** NAD(P)H $\tau_1$, **g)** NAD(P)H $\tau_2$, **h)** FAD $\alpha_1$, **i)** FAD $\tau_1$, and **j)** FAD $\tau_2$ for HDFs, EPCs, IMs, iPSCs and H9 ESCs (n = 459, 561, 990, 586, 35 respectively). **k)** Single-cell quantitative analysis of optical redox ratio for non-patterned (NP) and patterned (P) reprogramming cells (IMs and iPSCs). Data are presented as median with interquartile range for each cell type. Statistical significance was determined by one-way analysis of variance (ANOVA) using the Kruskal-Wallis test for multiple comparisons; ns = $p \geq 0.05$, * for $p < 0.05$, ** for $p < 0.01$, *** for $p < 0.001$, **** for $p < 0.0001$).

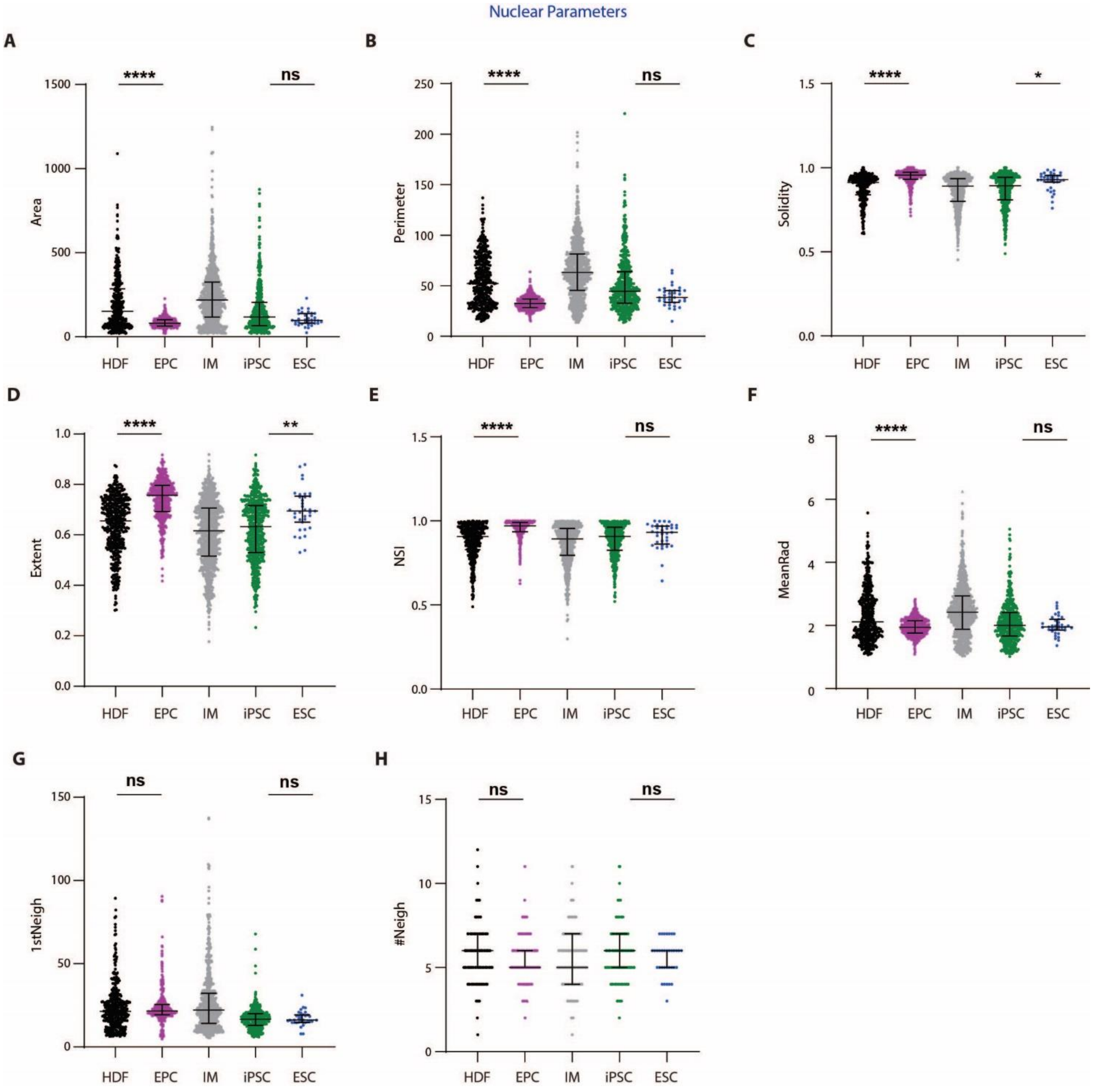


**Fig. S3. Nuclear parameter changes during reprogramming of EPCs.** Single-cell quantitative analysis of **a)** Area, **b)** Perimeter, **c)** Solidity, **d)** Extent, **e)** Nuclear Shape Index (NSI), **f)** Mean Radius **(**MeanRad), **g)** Distance to closest neighbor **(**1stNeigh), and **h)** Number of neighbors (#Neigh) for HDFs, EPCs, IMs, iPSCs and H9 ESCs (n = 459, 561, 990, 586, 35 respectively). Data are presented as median with interquartile range for each cell type. Statistical significance was determined by one-way analysis of variance (ANOVA) using the Kruskal-Wallis test for multiple comparisons; ns = $p \geq 0.05$, * for $p < 0.05$, ** for $p < 0.01$, *** for $p < 0.001$, **** for $p < 0.0001$).

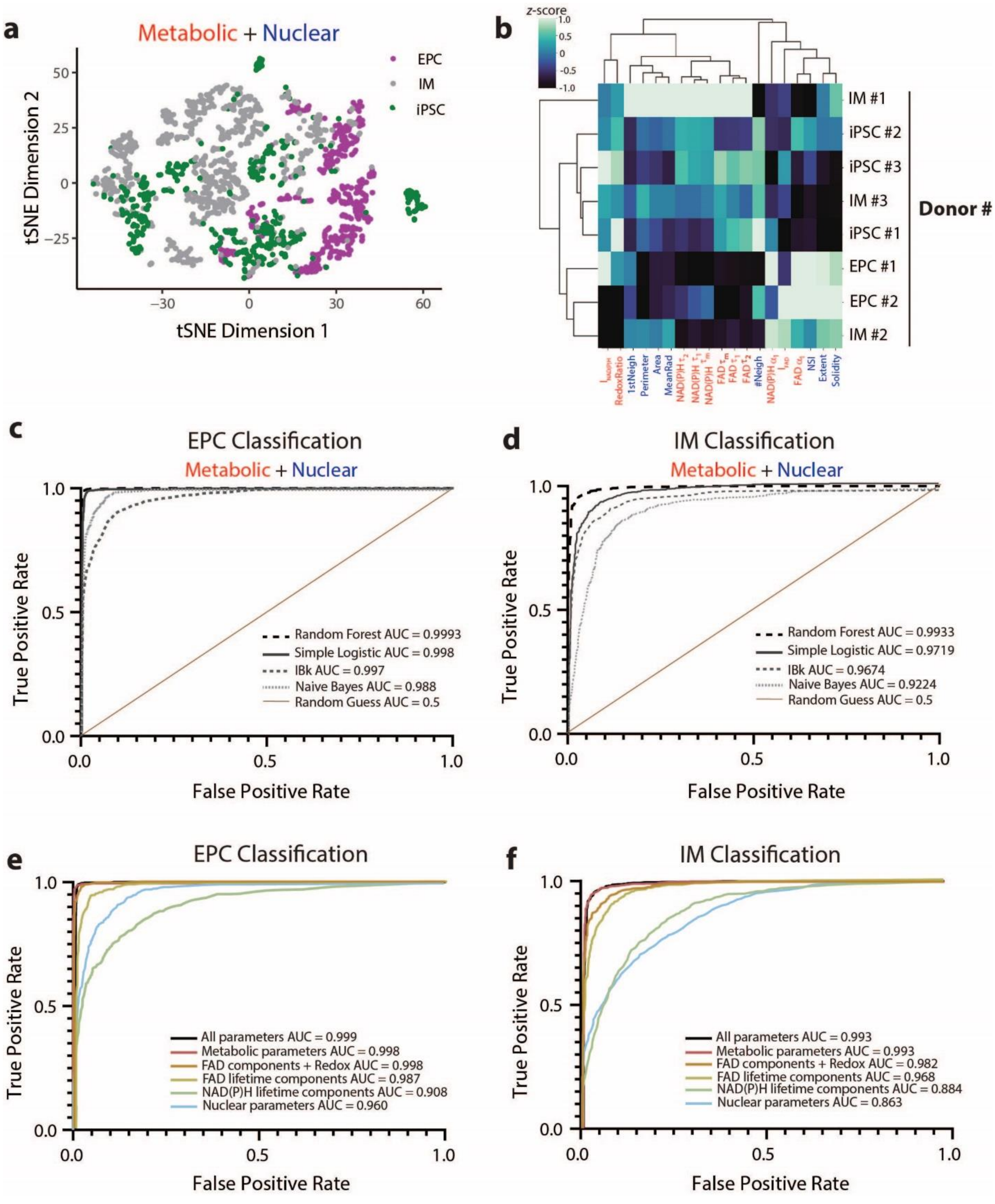


**Fig. S4. OMI enables accurate classification of EPCs, IMs, and iPSCs. a)** t-distributed stochastic neighbor embedding (t-SNE) dimensionality reduction was performed on all 11 metabolic and 8 nuclear parameters for each cell, projected onto 2D space, and shows poor separation of different cell types (EPCs, IMs, and iPSCs). Each color corresponds to a different cell type. Data are from three different donors. Each dot represents a single cell, and n =561, 990, and 586 cells for EPCs, IMs, and iPSCs, respectively. **b)** Heat map of $z$-scores ($z$-score = $(\mu_{observed}-\mu_{row})/\sigma_{row}$, where $\mu_{observed}$ is the mean value of each parameter for a cell type; $\mu_{row}$ is the mean value of each parameter for all cells together, and $\sigma_{row}$ is the standard deviation of each parameter across all cells. ) of metabolic and nuclear parameters; each row is the mean data aggregating all cells from a single donor and cell type (EPCs, IMs, iPSCs); n = 3 biologically independent donors. ROC curves for **c)** EPCs and, **d)** IMs for different classifiers computed using all 11 metabolic and 8 nuclear parameters. AUC is provided for each classifier as indicated in the legend. ROC curves for **e)** EPCs and, **f)** IMs for different classifiers computed using different parameter combinations. AUC is provided for each parameter combination as indicated in the legend.

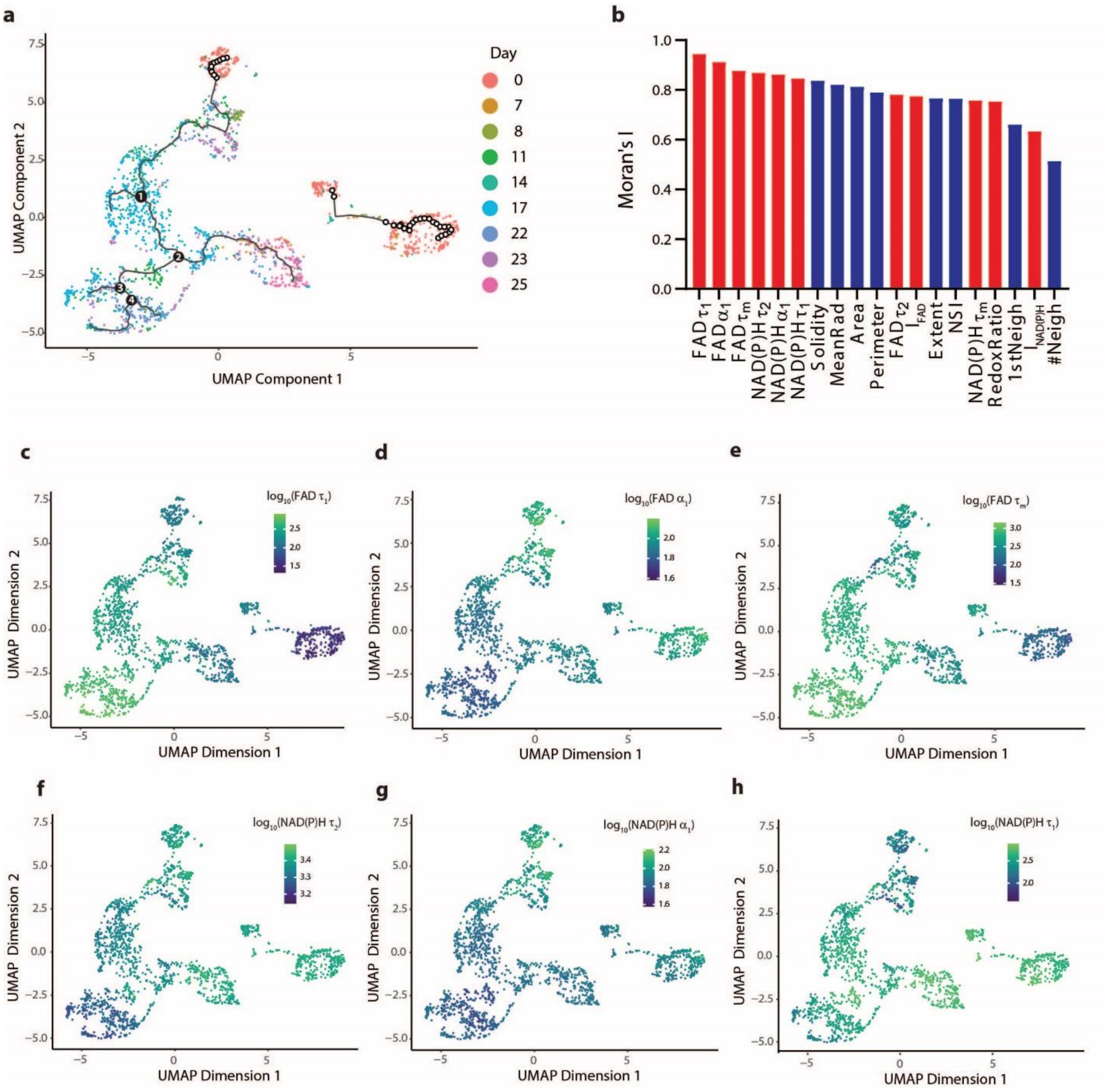


**Fig. S5. Metabolic and nuclear parameter changes along the reprogramming trajectory. a)** Trajectory of reprogramming EPCs constructed from the metabolic and nuclear parameters based on UMAP dimension reduction using Monocle, colored by day of EPC reprogramming. **b)** Metabolic and nuclear parameters ranked by their Moran's I on the construction of single-cell reprogramming trajectory. Moran's I value of 1 means that nearby cells will have perfect correlation, 0 represents no correlation, and -1 means that neighboring cells will be anticorrelated) UMAP plots based on Figure 3a highlighting the change of expression of top-six metabolic parameters **c)** FAD $\tau_1$, **d)** FAD $\alpha_1$, **e)** FAD $\tau_m$, **f)** NAD(P)H $\tau_2$, **g)** NAD(P)H $\alpha_1$ and, **h)** NAD(P)H $\tau_2$ during reprogramming.

**Supplementary files**

**Data S1. Metabolic and nuclear parameters of EPCs, IMs and iPSCs at different timepoints during reprogramming.**

**Data S2. Source data files for running the Monocle algorithm in R to generate single-cell reprogramming trajectories.**

**Code S1. R code used to generate single-cell reprogramming trajectories (Fig. 3.) using 11 metabolic and 8 nuclear parameters.**
